# Combining sample expansion and light sheet microscopy for the volumetric imaging of virus-infected cells with optical super-resolution

**DOI:** 10.1101/2020.04.10.035378

**Authors:** Luca Mascheroni, Katharina M. Scherer, James D. Manton, Edward Ward, Oliver Dibben, Clemens F. Kaminski

## Abstract

Expansion microscopy is a sample preparation technique that enables the optical imaging of biological specimens at super-resolution owing to their physical magnification, which is achieved through water-absorbing polymers. The technique uses readily available chemicals and does not require sophisticated equipment, thus offering super-resolution to laboratories that are not microscopy-specialised. Here we present a protocol combining sample expansion with light sheet microscopy to generate high-contrast, high-resolution 3D reconstructions of whole virus-infected cells. The results are superior to those achievable with comparable imaging modalities and reveal details of the infection cycle that are not discernible before expansion. An image resolution of approximately 95 nm could be achieved in samples labelled in 3 colours. We clearly resolve the concentration of viral nucleoprotein on the surface of vesicular structures within the cell and their positioning relative to cellular organelles. We provide detailed guidance and a video protocol for the optimal application of the method and demonstrate its potential to study virus-host cell interactions.

## 1. Introduction

Expansion microscopy is a technique that relies on the physical magnification of biological samples in order to visualise details that are spaced more closely than the diffraction limit of light (∼300 nm).^1 2^ The physical expansion is achieved by embedding fixed specimens in a polymer matrix that can absorb water, forming a so-called hydrogel. This approach generates volumetrically isotropic expansion and allows the bypassing of the diffraction barrier of light microscopy without any need for sophisticated instruments, enabling laboratories that possess standard fluorescence microscopes to image their samples at super-resolution. Despite the great potential of expansion microscopy, the physical nature of the expanded sample places some limitations on its use. Firstly, the hydrogels are challenging to image using conventional microscopes: such samples are bulky and mechanically unstable, making mounting and long-term imaging more challenging compared to conventional samples (for example, cells adherent on a glass slide). Secondly, when the hydrogel is placed in a dish and imaged through the bottom, the ‘vertical’ expansion of the sample hinders the imaging of the whole volume of the specimen using high numerical aperture (NA) objectives, which usually have short working distances. Moreover, the use of oil-immersion objectives generates a refractive index mismatch with the water-based hydrogels, which causes optical aberrations. Finally, the expansion process dilutes the fluorophore concentration, decreasing the fluorescence intensity up to a hundredfold, which hinders the imaging of expanded samples with microscopes incapable of collecting a large number of photons or those requiring very high fluorescent signals.

Expanded samples have so far been imaged mainly using confocal microscopes.^2^ However, a confocal microscope would not be optimal for the imaging of expanded samples: the weak fluorescence intensity of the expanded specimens is best recorded with a setup that is more photon-efficient than a confocal microscope. Light sheet microscopy is such a technique. Nonetheless, reports on the combination of light sheet microscopy with sample expansion have been limited so far.^3 4 5 6 7^

In light sheet microscopy, the optical pathways of excitation and detection are geometrically decoupled such that the sample is illuminated with a thin sheet of laser light, and detection is performed along an axis orthogonal to the illumination. This separation of excitation and detection light paths maximises detection efficiency by minimising out-of-focus fluorescence. The axial confinement of the excitation results furthermore in dramatically reduced photobleaching. For detection, fast cameras with high quantum efficiencies can be used, and thus imaging speed can be increased by orders of magnitude compared to point-scanning techniques.

The principle of light sheet microscopy was developed more than a century ago by Siedentopf and Zsigmondy, then termed ‘ultramicroscopy’.^8^ The method was rediscovered by developmental biologists at the beginning of the 21st century^9^ and its popularity has increased ever since. The high spatiotemporal resolution of light sheet microscopy was impressively demonstrated for high-speed imaging of embryonal development,^9^ neural activity,^10^ cardiac dynamics,^11 12^ and physiologically representative subcellular imaging.^13^ Despite all the advantages, a complication that regularly arises in light sheet microscopy concerns the mounting of the samples. The placement of samples on the microscope for imaging has to be compatible with the two-objective configuration of the method and solutions often need to be customised for the specific experimental configuration in use. Nowadays, commercial setups are available, and a lot of effort is still invested in the technical development of designs that are simpler to use and that improve the versatility and utility of the method.^14 15^

In this work, we study the suitability of expansion microscopy and light sheet microscopy for the imaging of virus-infected samples using human A549 cells infected with live attenuated influenza vaccine (LAIV), a modified low-virulence variant of the influenza A virus that is the basis of flu vaccine formulations and sold under the names Fluenz (USA and Canada) and FluMist (Europe).^16 17^ We show that expansion microscopy is particularly well suited to study host-cell vaccine interactions helping to understand underlying biomolecular mechanisms of the vaccine. Optical imaging at super-resolution is required in order to dissect the interplay of viral proteins and substructures with cell organelles, which often have dimensions comparable to, or smaller than, the diffraction limit of light (∼300 nm). However, imaging entire infected cells in multiple colours to build a full 3D picture of the infected cell is challenging using alternative super-resolution techniques, like *d*STORM^18^ and STED,^19^ due to their long acquisition times and proneness to photobleaching.

Here, we first describe the expansion of infected cells using a published expansion microscopy protocol. We then explain the imaging of the infected expanded cells on a light sheet microscope, and for comparison, also on widefield and confocal microscopes. By combining expansion microscopy with light sheet microscopy, we demonstrate how high-contrast 3D models of whole LAIV-infected cells can be easily reconstructed at super-resolution in three colour channels, highlighting the viral nucleoprotein located in cytosolic vesicles. These vesicles are essential for the transport of vRNA (viral RNA)-nucleoprotein complexes to the plasma membrane, where the mature virions form, and are also thought to mediate vRNA-vRNA interactions in the cytoplasm, which according to recent studies are crucial for virus assembly.^20^ The vesicles were not resolved without expansion, but due to expansion could be characterised for their size and cellular distribution. Finally, we present a detailed video protocol on the mounting and imaging of expanded samples using a light sheet microscope. The aim is to make this technique available to the wider community and so that the power of light sheet and expansion microscopy can be harnessed for addressing questions on virus research specifically, but also in cell biology more generally, for problems that require the study of detail within cellular volumes at sub-wavelength resolution.

## 2. Materials and methods

### 2.1 Chemicals

Methanol-free formaldehyde was purchased from Thermo Fisher Scientific; the ampoules were used immediately after opening and any leftover formaldehyde discarded. All chemicals used for sample expansion (glutaraldehyde 50% in water, sodium acrylate, N, N’-methylenbisacrylamide, acrylamide, Proteinase K) were purchased from Sigma Aldrich and used as received.

### 2.2 Antibodies

Mouse anti-influenza A nucleoprotein (ab20343) and rabbit anti-beta tubulin (ab6046) primary antibodies were purchased from Abcam. Detection was via polyclonal goat secondary antibodies: an ATTO647N-conjugated anti-mouse antibody was purchased from Sigma Aldrich, while an AlexaFluor488 (AF488)-conjugated anti-rabbit antibody was purchased from Invitrogen.

### 2.3 Virus

The live attenuated influenza vaccine (LAIV, 9.2 TCID50/mL) derived from the wildtype Influenza A strain A/New Caledonia/20/99 was provided by AstraZeneca (Speke, Liverpool).

### 2.4 Cell cultures

A549 cells were purchased from the European Collection of Authenticated Cell Cultures (ECACC). The cell line was cultured at 37°C and 5% CO_2_ in Dulbecco-modified MEM (Sigma Aldrich) supplemented with 10% heat-inactivated foetal bovine serum (Gibco), antibiotics/antimycotics (100 units/mL penicillin, 100 µg/mL streptomycin, 0.025 µg/mL Gibco Amphotericin B, Gibco) and 2 mM L-glutamine (GlutaMAX, Gibco). Cultures at ∼80% confluency were routinely split into T-75 polystyrene flasks.

### 2.5 Infection of A549 cells with LAIV

A549 cells were plated onto 13 mm round coverslips in 4-well plates at 60,000 cells per well, 16 hours before infection. The next day, cells were infected with LAIV at 10 PFU per cell. After one hour of incubation at 37°C and 5% CO_2_, the medium was exchanged with fresh new medium. Cells were fixed 9 hours post infection (hpi), permeabilised and labelled with antibodies according to procedures described below.

### 2.6 Immunostaining

Infected cells were fixed by incubation with 4% methanol-free formaldehyde and 0.1% glutaraldehyde in PBS for 15 minutes at room temperature, washed three times with PBS and then permeabilized by incubation with a 0.25% solution of Triton X-100 in PBS for 10 minutes. Unspecific binding was blocked by incubating with 10% goat serum in PBS for 30 minutes at room temperature. Without washing, the samples were incubated with the primary antibody, diluted 1:200 in PBS containing 2% BSA (bovine serum albumin) for 1 hour at room temperature. After three washes in PBS, the samples were incubated with the secondary antibody, diluted 1:400 in PBS containing 2% BSA, for an hour at room temperature in the dark. Samples were then washed 3 times with PBS. Samples that were not meant for expansion microscopy were counterstained with DAPI nuclear dye (10 µg/mL in PBS for 10 minutes at room temperature). Finally, the coverslips were mounted on glass slides using a Mowiol-based mounting medium. Alternatively, samples were expanded using the expansion microscopy protocol detailed below.

### 2.7 Expansion and imaging of samples

The expansion of samples was achieved following a published protocol.^21^ Nuclear staining was performed after the first round of expansion, using DAPI, 10 µg/mL in water for 20 minutes. In order to image the expanded gels on the widefield and confocal microscopes, they were cut using a glass coverslip as a knife to fit in glass-bottom Petri dishes, which were pre-coated with poly-L-lysine (0.02% in water for 30 minutes). Alternatively, the gels were imaged with the light sheet microscope, by cutting a strip of gel with cells facing up, which was then glued on a 24×50 mm glass coverslips using cyanoacrylate-based super-glue (Henkel). The slide was left to cure for two minutes and then placed in an imaging chamber and filled with milli-Q water. This process is documented in detail in Supporting Video 1.

### 2.8 Microscopes

The light sheet microscope was home built using the inverted selective plane illumination microscopy (iSPIM)^22^ design for which parts were purchased from Applied Scientific Instrumentation (ASI, Eugene, USA) including controller (TG8_BASIC), scanner unit (MM-SCAN_1.2), right-angle objective mounting (SPIM-K2), stage (MS-2K-SPIM) with motorized Z support (100 mm travel range, Dual-LS-100-FTP) and a filter wheel (FW-1000-8). All components were controlled by Micro-Manager, using the diSPIM plugin from ASI. The setup was equipped with a 0.3 NA excitation objective (Nikon 10x, 3.5 mm working distance) and a 0.9 NA detection objective (Zeiss, W Plan-Apochromat 63x, 2.1 mm working distance) to maximise spatial resolution and fluorescence signal collection. Lasers were fibre-coupled into the scanner unit. An sCMOS camera (ORCA-Flash 4.0, Hamamatsu, Hamamatsu-City, Japan) was used to capture fluorescence. DAPI was excited with 445 nm (OBIS445-75 LX), AlexaFluor488 with 488 nm (OBIS488-150 LS) and ATTO647N with 647 nm (OBIS647-120 LX). For fluorescence detection, respective emission filters were used (DAPI: BrightLineFF01-474/27, AF488: BrightLineFF01-540/50, ATTO647N: BrightLineFF0-680/42). Filters were purchased from Semrock (New York, USA).

The widefield microscope was home built and parts were purchased from the following suppliers: frame (IX83, Olympus, Tokyo, Japan), stage (Prior, Fulbourn, UK), Z drift compensator (IX3-ZDC2, Olympus, Tokyo, Japan), plasma light source (HPLS343, Thorlabs, Newton, USA), and camera (Clara interline CCD camera, Andor, Belfast, UK) of the custom-built wide-field microscope were controlled by Micro-Manager. Respective filter cubes for DAPI (excitation 350 nm, dichroic mirror 353 nm, emission 460 nm), AF488 (excitation 500 nm, dichroic mirror 515 nm, emission 535 nm) and ATTO647N (excitation 635 nm, dichroic mirror 660 nm, emission 680 nm) were used for selecting excitation and detecting fluorescence. The images were acquired with an Olympus PlanApoU 60x/1.42 oil objective lens.

The confocal microscope was a commercial Leica TCS SP5. The following excitation lasers were used: 405 nm (DAPI), 488 nm (AF488), 647 nm (ATTO647N). The following detectors were used: DAPI 420-450 nm; AlexaFluor488 500-600 nm; ATTO647N 660-800 nm. The images were acquired with a 63x/1.40 oil objective lens.

### 2.9 Data analysis

Vesicle sizes present in expanded samples that were imaged using light sheet microscopy were analysed using ImageJ. Data were plotted in GraphPad Prism (GraphPad Software, US).

## 3. Results and discussion

### 3.1 Infection, staining and expansion of cells

A549 cells, from human alveolar carcinoma, are a common model for the study of LAIV infection and replication.^17^ We incubated the cells with the LAIV particles at 10 PFU for one hour, then we exchanged the medium and let the infection cycle progress until fixation. The LAIV virions are not fluorescent, therefore, immunostaining of the viral proteins is necessary in order to study the infection progression using fluorescence microscopy. Here, we stained the LAIV nucleoprotein (NP), a structural protein that packs the viral RNA inside of the virus.^23^ Additionally, we stained the cell microtubules and the cell nuclei, in order to study the interaction between the viral particles and cell organelles. A picture of the stained infected cells, not expanded and imaged on a confocal microscope, is shown in Figure 1. The resolution of the non-expanded images is enough to localise the viral nucleoprotein (NP) in the cell cytosol. However, the image resolution is too low to clarify the exact NP localization, form, and interaction with other cellular structures.

**Figure 1.**
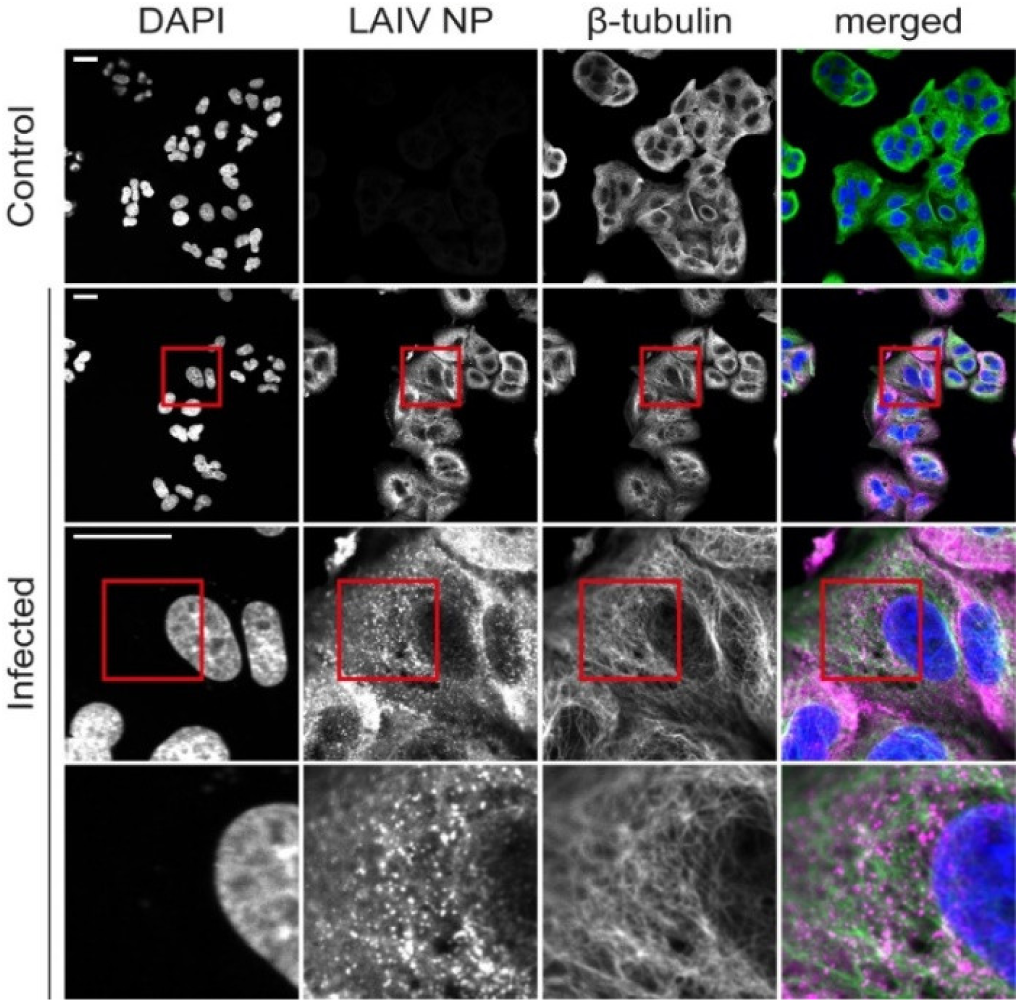
Non-expanded A549 cells infected with live attenuated influenza vaccine (LAIV). Cells were fixed 9 hours post infection and images recorded on a confocal microscope. Immunostaining was used to fluorescently label LAIV nucleoprotein (NP, magenta) and microtubules (green) whereas DAPI staining was used for labelling the nuclei (blue). Scalebar 25 µm. Size of images in bottom row 25×25 µm.

After immunostaining, the cells were expanded using a published expansion microscopy protocol.^21^ Briefly, expansion microscopy works by synthesising a polymer matrix *in situ*, which cross-links the protein structures of the sample. The sample, now embedded within the gel, is then enzymatically digested by proteases in order to cleave the cells’ rigid structures, such as the cytoskeleton. Without digestion, the gelled sample would not expand; however, the linking to the gel matrix guarantees that the cleaved proteins are not lost and that they keep their relative positions. Finally, the gel is placed in deionised water to expand. The expansion process spatially separates the fluorophores that are spaced more closely than the scale of the microscope resolution, hence increasing the level of detail in the final image. A schematic representation of the steps of the expansion microscopy protocol is shown in Supporting Figure 1.

### 3.2 Mounting expanded samples for microscopy

The expanded gels are unconventional imaging samples: since they mainly consist of water, they are very fragile and unsteady. In order to image the gelled samples on inverted confocal or widefield microscopes, we cut them to fit into glass-bottom Petri dishes with cells facing down (Figure 2A, left). One issue we encountered while imaging the gels in this configuration is their wobbling and drifting during the image acquisition. To minimise this issue, we pre-coated the glass bottom surfaces of the dishes with poly-L-lysine, which improved gel adherence. However, this is not always sufficient to keep the gels still, especially during long-term imaging. The use of cyanoacrylate-based glue has been suggested to keep the gels in place,^24^ although here this would require the glue to be placed in direct contact with the cell-containing side of the sample, leading to potential deterioration of embedded biological structures. Moreover, the use of glue means that the glass bottom cannot be reused.

**Figure 2.**
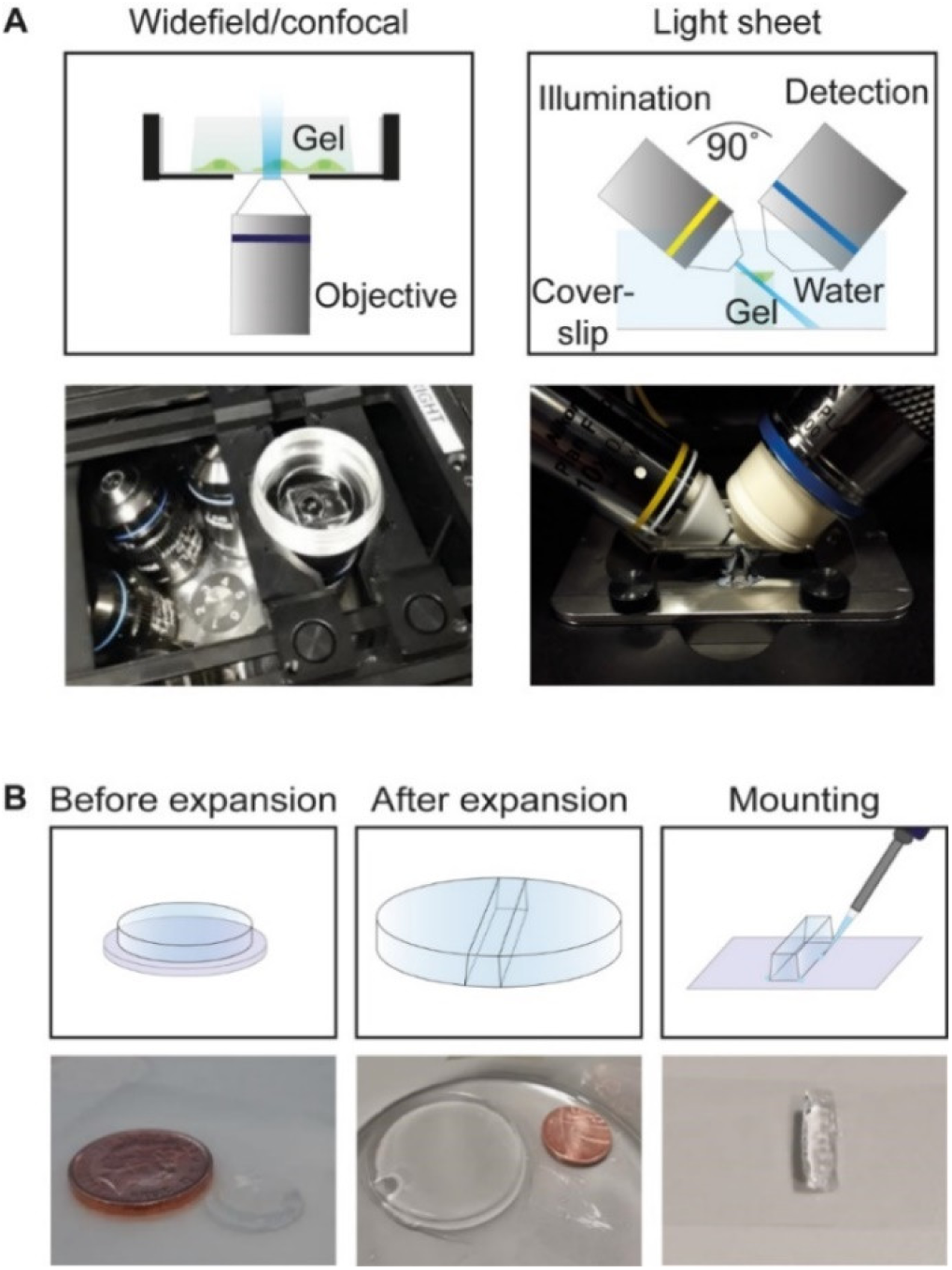
(A) Schematic representation of gel mounting for widefield/confocal microscopy (left) on an inverted microscope and for light sheet microscopy on a custom-built iSPIM setup (right). (B) Step-by-step procedure for light sheet microscopy.

In order to image the expanded samples using a light sheet microscope, a small gel strip was cut and attached to a glass slide (24×50 mm) using super-glue, with cells facing up (Figure 2A, right). In this configuration, the glue was not in direct contact with the cells. The procedure was effective in preventing problems associated with gel wobbling or drift. Cutting the gel into a strip permits it to be placed in the small space between the two light sheet objectives; this configuration enables the imaging of the whole depth of the sample. Moreover, the light sheet microscope is equipped with water-dipping objectives, which eliminates optical aberrations due to refractive index mismatches with the (water-based) gels. On the other hand, the confocal and widefield microscopes that we used for this study were optimised for the imaging of fixed samples and were equipped with high NA oil objectives for the purpose of high light collection efficiency. They are however not ideal for imaging gelled samples, whose optical properties resemble more those of living samples than fixed ones. A step-by-step procedure for the mounting of expanded samples on a light sheet microscope is depicted in Figure 2B. A detailed video protocol of this procedure is presented in Supporting Video 1, where we show how to cut the gelled sample and glue it to the glass slide of the imaging chamber of the light sheet microscope.

### 3.3 Comparison of imaging modalities

We compared the performance of light sheet microscopy of expanded samples with two commonly used conventional fluorescence microscopy techniques, widefield and confocal laser scanning microscopy (CLSM). The working principles of all three techniques are illustrated in Figure 3A. When imaging a sample using a widefield microscope, the whole fluorescent specimen is excited, which results in considerable out-of-focus light derived from fluorophores that lie outside of the focal plane, producing high background signals and low image contrast. A confocal laser scanning microscope mitigates this problem by using a point-source for illumination and pinholes to filter out out-of-focus fluorescence. As a result, CLSM features better image contrast compared to widefield microscopy for thick samples. However, the requirement for scanning the point sequentially across the sample decreases the acquisition speed significantly. A light sheet microscope combines the advantages of a confocal laser scanning and a widefield microscope: here the sample is illuminated with a thin sheet of light that excites only those fluorophores that lie within the vicinity of depth of field of the detection objective. Thus, signal is only generated from the illuminated fluorophores and those in out-of-focus planes are not excited and therefore do not contribute to image blur. Speed is high due to the parallel detection of the widefield signal by all pixels of a camera.

**Figure 3.**
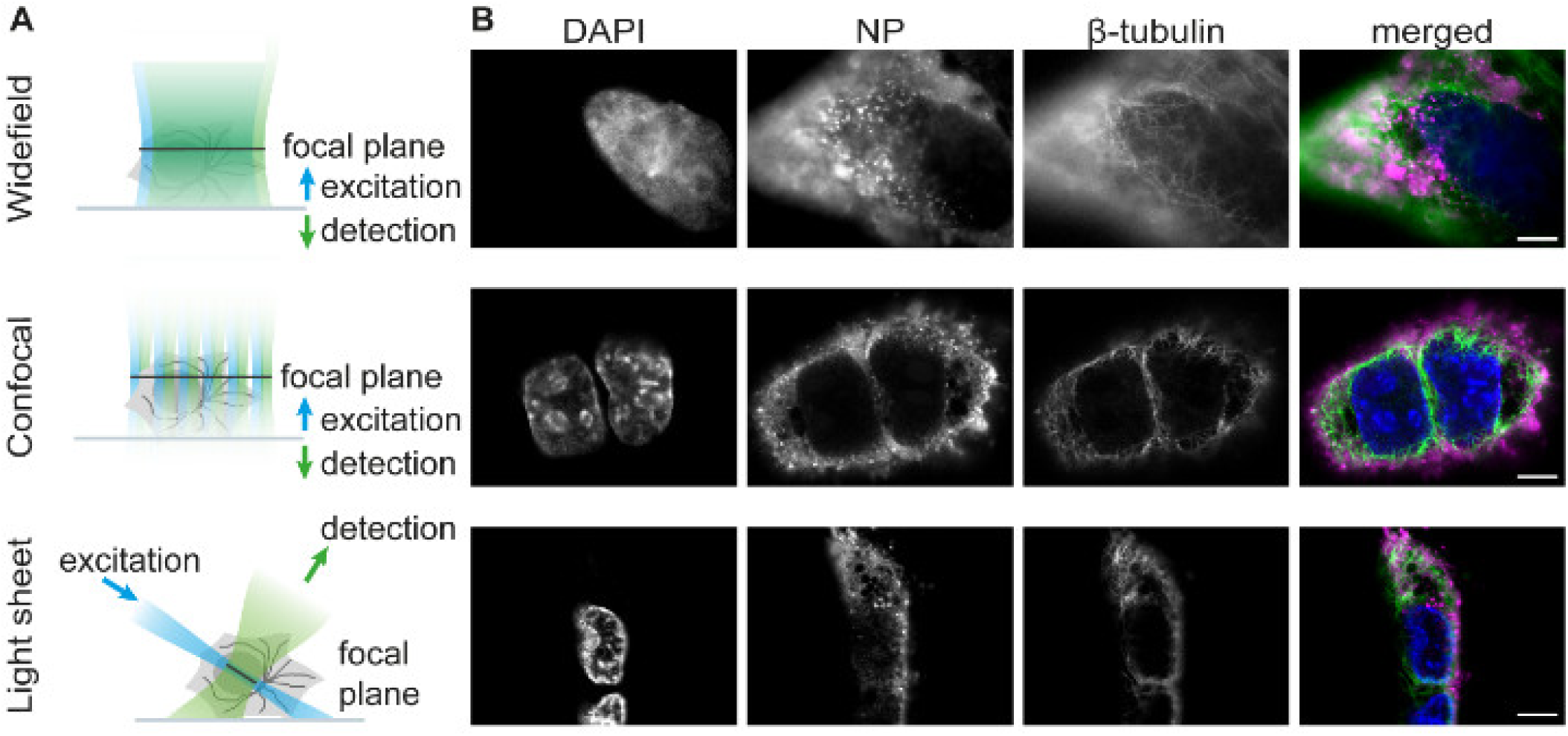
(A) Comparison of three different fluorescence microscopy techniques for imaging of expanded samples. (B) Expanded A549 cells infected with live attenuated influenza vaccine (LAIV). Cells were fixed 9 hours post infection, immunostained, expanded and imaged using widefield, confocal or light sheet microscopy. Merged images show LAIV nucleoprotein in magenta, microtubules in green and cell nuclei in blue. Scale bar 25 µm.

Using all three microscopy techniques, we imaged A549 cells that were infected with live attenuated influenza vaccine (LAIV), fixed 9 hours post-infection, immunostained and expanded as described in Section 3.1. Example images featuring the cell nucleus, the viral nucleoprotein (NP) and microtubules are displayed in Figure 3B. As expected, we observed the lowest amount of bleaching with the light sheet microscope. The acquisition of a stack composed of 200 frames took roughly 30 seconds, using an exposure time of 50 ms; after acquiring one stack, we did not notice any evident sample bleaching. On the widefield microscope, photobleaching after an acquisition of a comparable volumetric image stack was also low. CLSM, however, resulted in substantial photobleaching due to the high laser power used (up to 80 mW) to boost the fluorescent signal. Furthermore, the speed of CLSM was also significantly reduced compared to the other methods because of its point scanning nature. Acquisition for a single frame in three colours took roughly five minutes by CLSM (using a line scanning frequency of 10 Hz and a 2048×2048 pixel field of view), while it took roughly 5 seconds on the widefield setup (200 ms exposure) and less than a second on the light sheet microscope (50 ms exposure). This long acquisition time, combined with the substantial photobleaching noticed, led us to rule out CLSM as a valid way of scanning our sample volumetrically.

We note that the microscope pinhole was opened to correspond to 3.7 Airy units in size (in contrast to the pre-set value 1.0 Airy units). This was done to boost the weak fluorescence signal, but came at the cost of decreased image contrast. In principle, similar (or better) contrast to light sheet microscopy can be achieved with CLSM via the use of small pinholes, but this is possible only for samples of high brightness. The reduced sensitivity of the confocal system reflects fundamental differences in the detection of the fluorescent signal.

To assess the image quality produced by the different imaging modalities, we applied a Fourier spectral power analysis similar to the method proposed by Demmerle et al.^25^ Representative images of LAIV NP were selected and Fourier transformed, then radially averaged to produce a spectral power plot (Figure 4A). This shows that the light sheet images have better spectral power at all spatial frequencies, denoting superior contrast. Furthermore, the spatial frequency at which each curve crosses the noise floor, denoting the absolute resolution limit, is the same for both the confocal and light sheet case. This demonstrates that the effective resolution of both types of image is approximately 400 nm.

**Figure 4.**
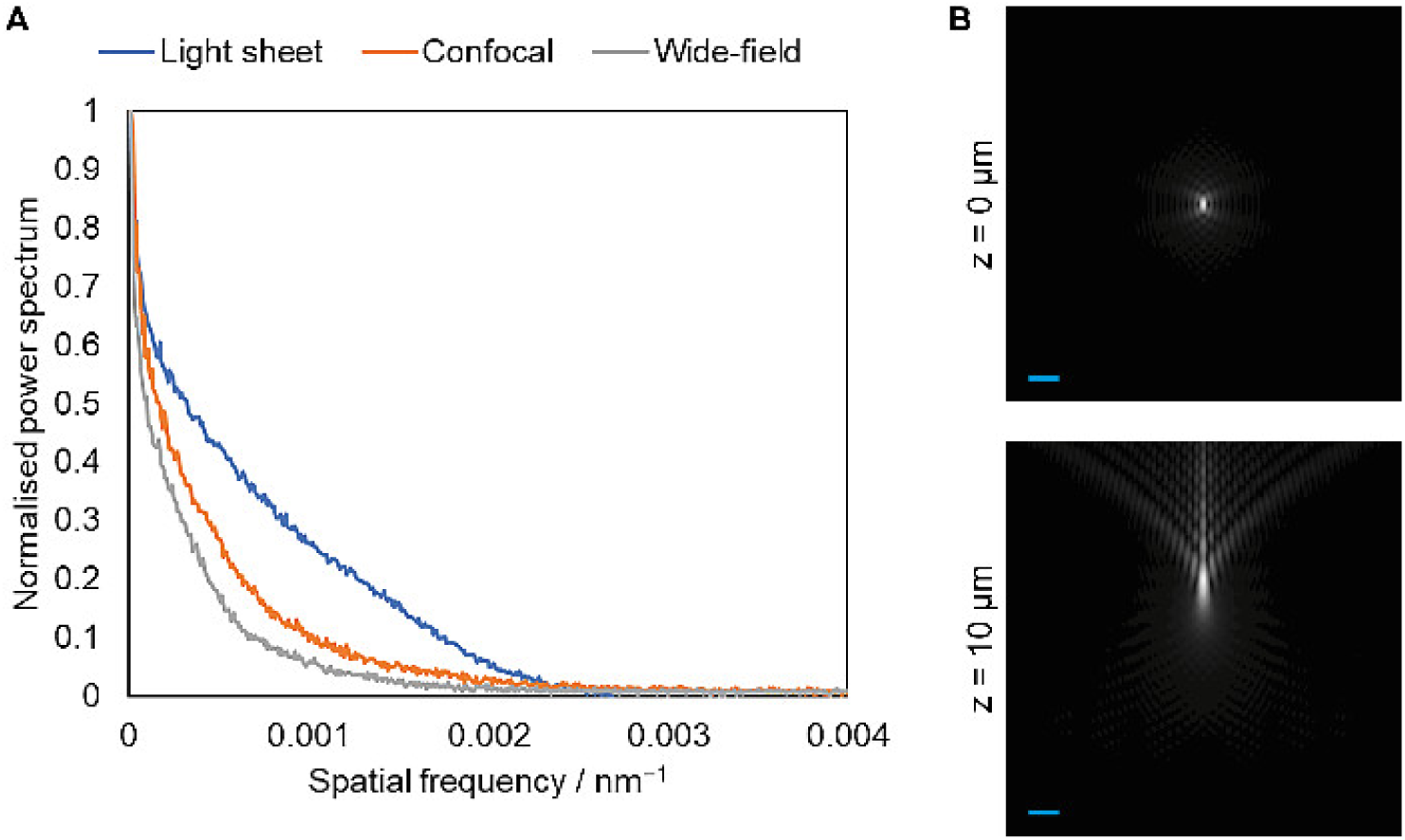
(A) Spectral power analysis of light sheet, confocal and wide-field images. No technique has appreciable spectral power beyond 0.0025 nm^−1^, corresponding to an ultimate optical resolution limit of 400 nm. (B) Simulated XZ slices of point spread functions for the oil immersion lenses used for wide-field and confocal imaging at expanded gel sample depths of 0 and 10 microns. A significant aberration is shown at only 10 micron depth, halfway through the depth of a typical expanded cell, due to the significant difference in refractive index between gel and oil immersion. Scale bar 1 µm.

This effective resolution limit is close to that predicted by the Abbe resolution criterion (λ / 2 NA) for the light sheet case in which the NA is 0.90 (375 nm), but is far from the theoretical resolution limit of 255 nm for the oil immersion lens used in the confocal imaging. We hypothesised that this was due to the aberrations induced by imaging at depth in a watery sample. Using a model for index-mismatched point spread functions^26^ and a vectorial diffraction simulation,^27^ we calculated theoretical point spread functions (PSF) for the oil immersion lens at different depths within the watery sample (Figure 4B). This showed that the PSF was already highly aberrated only 10 µm into the sample, corresponding to roughly half the thickness of several types of mammalian cells.

These simulated PSFs also showed significant ‘flare’ at depth, in which much of the intensity within the PSF is displaced away from the focal plane. This explains why the spectral power analysis shows the widefield image to have the worst image quality and resolution (∼500 nm): in contrast to the confocal case, there is no mechanism with which to exclude this out-of-focus light, leading to significant out-of-focus blur contamination.

In contrast to widefield and confocal microscopy stacks, the light sheet images require post-imaging processing, specifically a procedure called deskewing. This is necessary to correct for the angle at which the stage scans the sample through the focal plane of the detection lens (48° with respect to the detection axis) to reconstruct the pictures in a conventional geometry. The deskewing can be performed with an affine transformation where each image slice is computationally shifted (deskewed) to its proper position in the three-dimensional image volume. This procedure was performed using a customized ImageJ macro based on the TranformJ ImageJ plugin.^28 29^

### 3.4 Expansion microscopy highlights the vesicular structure of NP-containing compartments

By combining expansion and light sheet microscopy we could generate high-contrast 3D reconstructions of whole infected cells. In Figure 5A, we show maximum intensity projections at different magnifications obtained from a light sheet image z-stack after deskewing. Using the z-stacks acquired combining expansion microscopy and light sheet microscopy we could render 3D models of whole LAIV-infected A549 cells with high contrast, as shown in Figure 5A (fourth row) and Supporting Video 2. The resolution increase brought about by the sample expansion allowed us to demonstrate that the LAIV nucleoprotein localises at or in the membrane of small vesicular structures in the cell cytosol (Figure 5, third row). It is assumed that viral ribonucleoprotein complexes (vRNP) of influenza A virus, which are formed by nucleoprotein, viral RNA and other viral proteins, start to assemble in the cell cytoplasm at the membranes of recycling endosomes and are transported by those vesicles to sites of virion formation at the plasma membrane. ^30 31 32 33^ Taking the calculated expansion factor of 4.2 (Figure 5B) into account, we find that the vesicles possess a diameter of up to ∼500 nm (see size distribution in Figure 5C). The smallest vesicle diameter that we could resolve is ∼150 nm. In contrast, before expanding the sample we were not able to resolve the vesicular structure of the compartments occupied by the viral nucleoprotein (Figure 1). Interestingly, we find that the larger vesicles preferably occupy a space next to the nucleus where typically the Golgi apparatus is positioned, whereas the smaller vesicles are closer to the cell periphery. The space occupied by the larger vesicles is almost devoid of microtubules, and the vesicles do not seem to directly interact with the microtubules. This is interesting and in line with observations for influenza A virus, which suggest that the mode of transport for vRNP complexes can be microtubule-independent.^31^ Moreover, there was no identifiable DNA compaction inside the nucleus, which is typical of other viruses such as herpes.^34^ In the future, we aim to use this technique for studying the interplay between the viral proteins and the cell compartments in order to dissect the whole replication cycle of LAIV.

**Figure 5.**
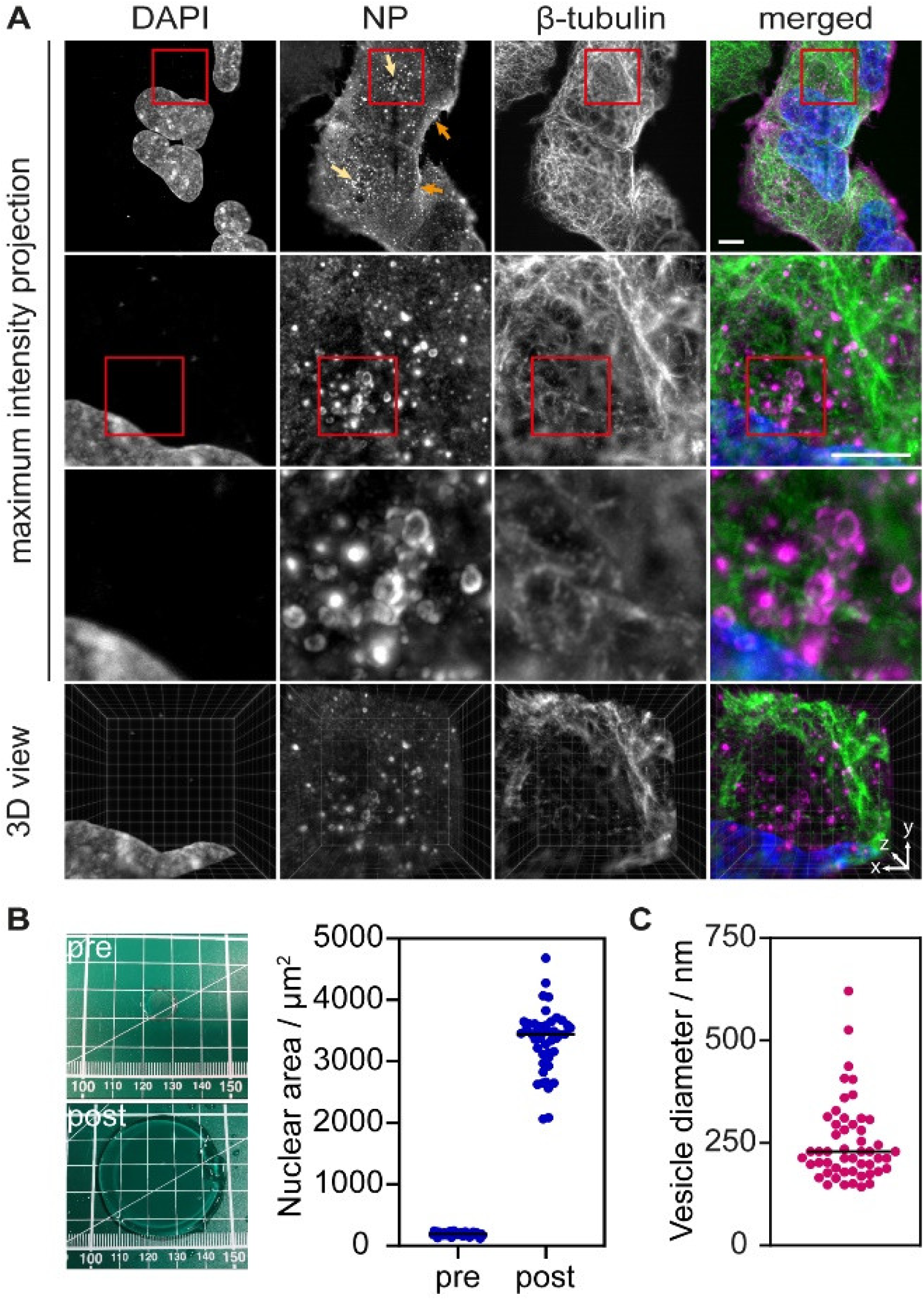
(A) Maximum intensity projections of expanded A549 cells infected with live attenuated influenza vaccine (LAIV). The expansion allows resolving the structure of the viral nucleoprotein NP, which forms vesicular structures in the cytosol before the virus egress. Cells were fixed 9 hours post infection, immunostained, expanded and imaged using a light sheet microscope. Cell nuclei are shown in blue, LAIV nucleoprotein (NP) in magenta and microtubules in green. Scale bar 20 μm. Images in the third row are 20 × 20 µm^2^. Fourth row shows a snapshot by the ClearVolume plugin for ImageJ^37^. Arrows in the first row NP image indicate small vesicles at the cell periphery (orange) and big vesicles at the perinuclear region (bright yellow). (B) A linear expansion factor of 4.2 (corresponding to a volumetric expansion factor of 74) was calculated by measuring the gel size before and after expansion (left panel), as well as by measuring the area of 50 cell nuclei before and after expansion (right panel). (C) The expansion factor was used to calculate the size of the NP-containing vesicles.

## 4. Conclusions

In this work, we imaged LAIV-infected A549 cells using a combination of expansion microscopy and light sheet microscopy, as well as confocal and widefield microscopy. The 3D rendering of whole infected cells was troublesome using data acquired by confocal microscopy, given the long acquisition time and intense photobleaching noticed in the gelled sample portion. The widefield microscope did not possess enough sectioning resolution and was characterised by low image contrast, thus, it could not produce cell reconstructions with a sufficient level of detail for our purpose. Using the light sheet microscope, instead, we were able to scan the whole specimen and deliver 3D renderings of whole infected cells with the highest level of image quality of the three techniques. We concluded that light sheet microscopy in combination with sample expansion is most suitable for detailed investigations of the interplay between the viral proteins and the cell organelles in whole cells.

In terms of sample mounting and imaging, light sheet microscopy poses challenges, owing to the gel nature of the sample. We include instructions on the procedure in a step-by-step video protocol and find that on mastering the method, the workflow for imaging with light sheet microscopy was faster than that for either widefield or confocal microscopy. While the acquisition of a stack composed of 200 frames took less than a minute using the light sheet microscope (50 ms exposure time), on the confocal system it took around 5 minutes or the acquisition of a single frame. We note that increasing the scanning speed or reducing the field of view would decrease the time of confocal imaging significantly, nonetheless, we are confident that a confocal system could not outperform a light sheet microscope as regards acquisition speed.

Expansion microscopy is a technique that has not yet been widely explored in virus research and has so far only been applied in two very recent studies on herpes virus simplex 1.^7 35^ However, we show that it provides unprecedented detail on the interaction and localisation of virus particles with subcellular organelles. We observe the nucleoprotein (NP) in the membrane of cytoplasmic vesicles which are up to 500 nm in size and larger in the perinuclear region compared to the cell periphery. From previous studies on influenza A it is known that cellular Rab11a-containing endosomes colocalize with vRNPs.^31^ Using our methodology, we can now characterise these vesicles in size and cellular distribution. This is important because these vesicles are essential for the transport of vRNPs to the plasma membrane where the mature virions form and are thought to mediate vRNA-vRNA interactions which according to recent studies are crucial for influenza virus assembly. ^36^

The advantages of combining expansion and light sheet microscopy were here demonstrated in a study of LAIV, but the method is, of course, applicable for studies of host cell biology in general. The continuous development of new protocols will allow the investigation of distinct events in the viral life cycle like entry, assembly and egress with high resolution since viral proteins, host cell proteins and even viral RNA can potentially be visualised at the same time.

## Acknowledgements

CFK acknowledges funding from the UK Engineering and Physical Sciences Research Council, EPSRC (grants EP/L015889/1 and EP/H018301/1), the Wellcome Trust (grants 3-3249/Z/16/Z and 089703/Z/09/Z) and the UK Medical Research Council, MRC (grants MR/K015850/1 and MR/K02292X/1), AstraZeneca, and Infinitus (China) Ltd. We also thank the Department of Chemical Engineering and Biotechnology of the University of Cambridge for providing the Leica TSC SP5 microscope and Francesca van Tartwjik for proofreading the manuscript.

## SUPPORTING INFORMATION

**Supporting Figure 1.**
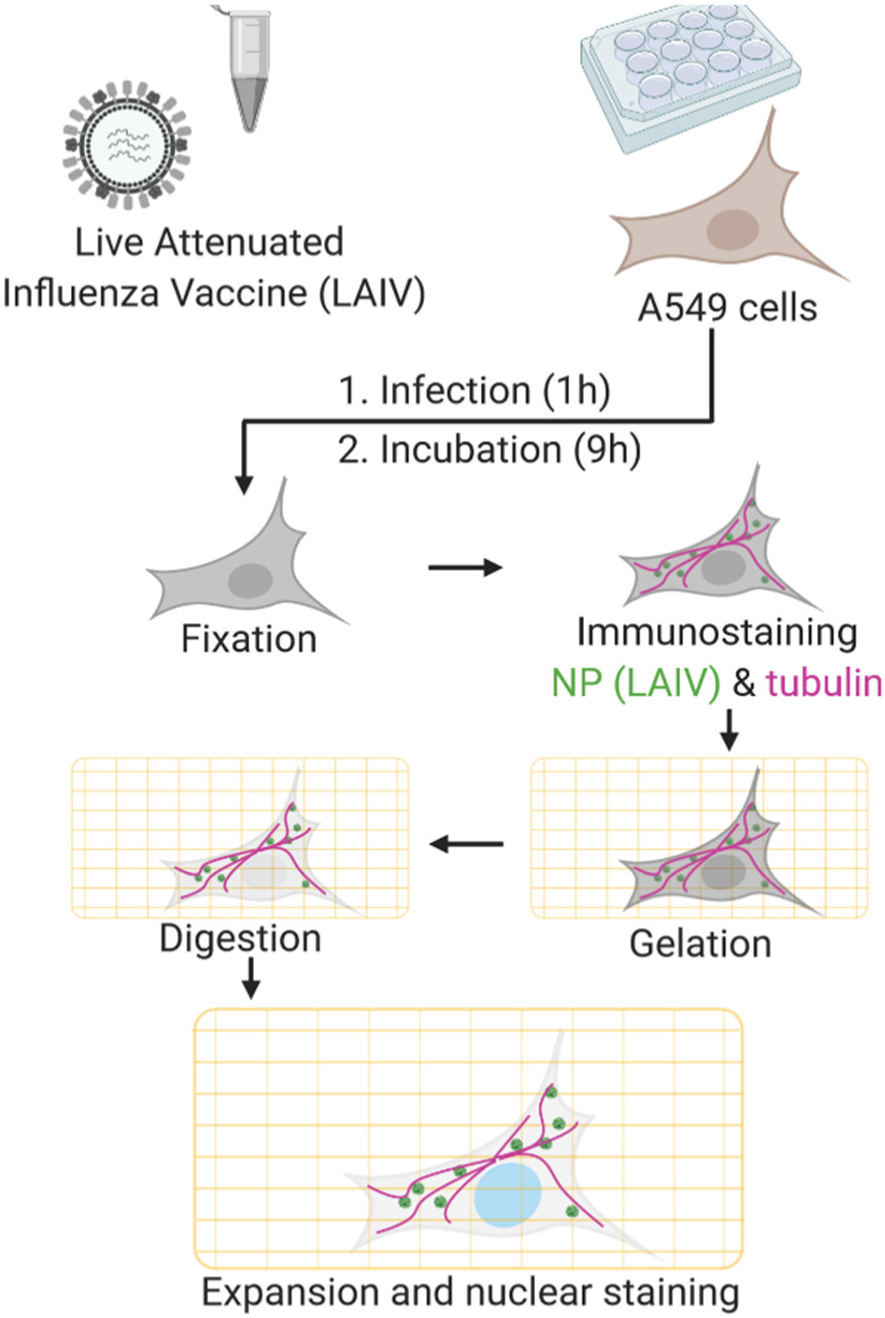
Schematic rendering of the infection with live attenuated influenza vaccine, immunostaining and expansion of A549 cells. Image created using BioRender.

**Supporting Figure 2.**
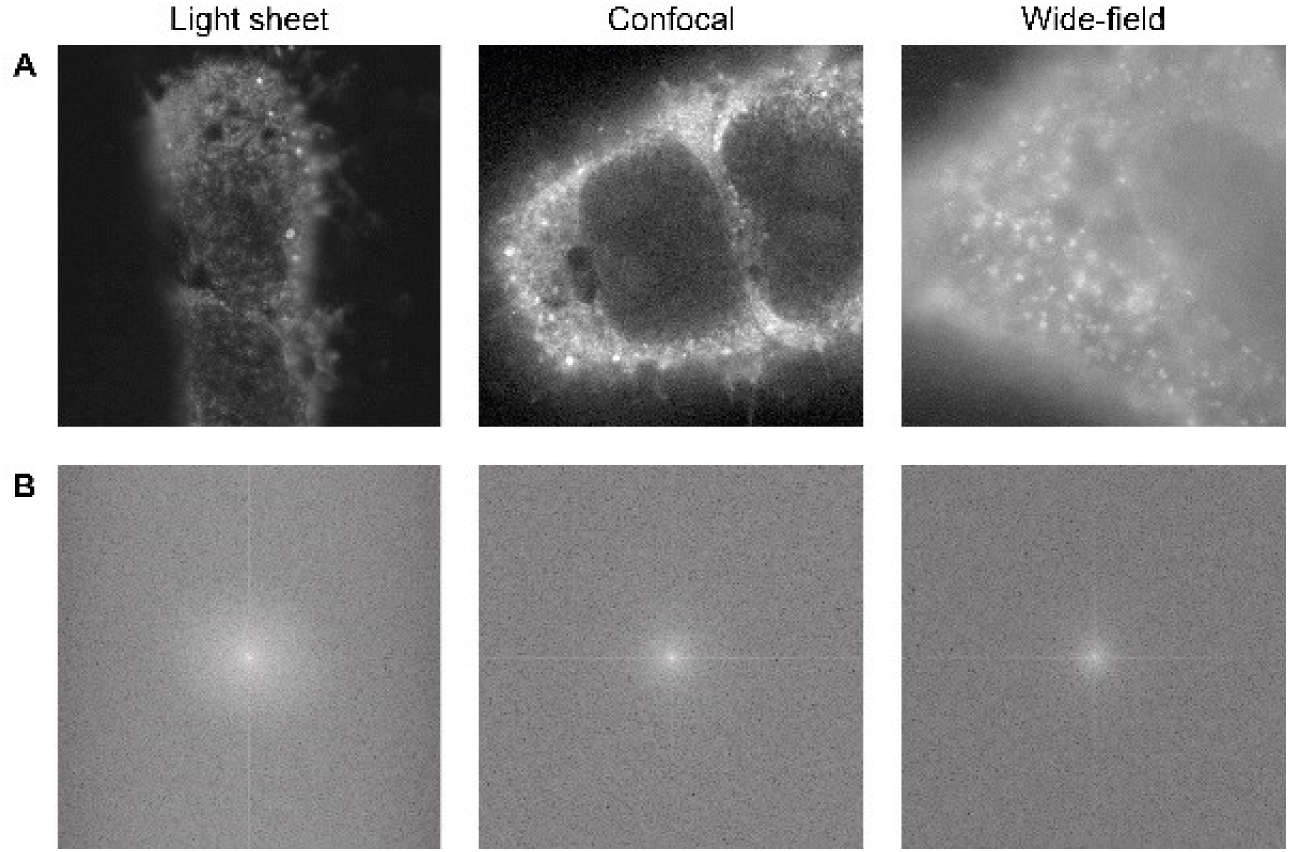
(A) Representative images of immunostained NP in expanded A549 cells infected with LAIV displayed with a gamma of 0.5. (B) Fourier power spectra of images in (A).

**Supporting Video 1.**
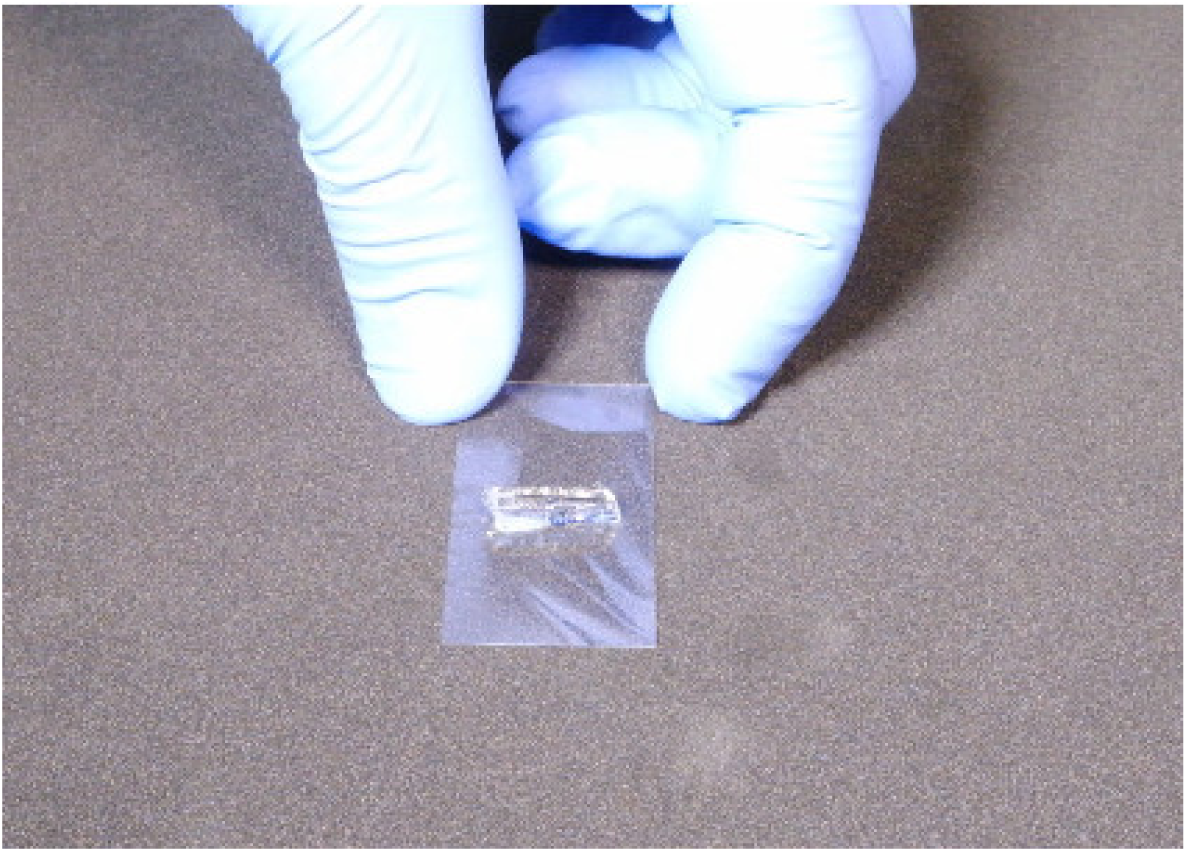
This video shows a detailed step-by-step procedure for the mounting of an expanded sample on a light sheet microscope.

**Supporting Video 2.**
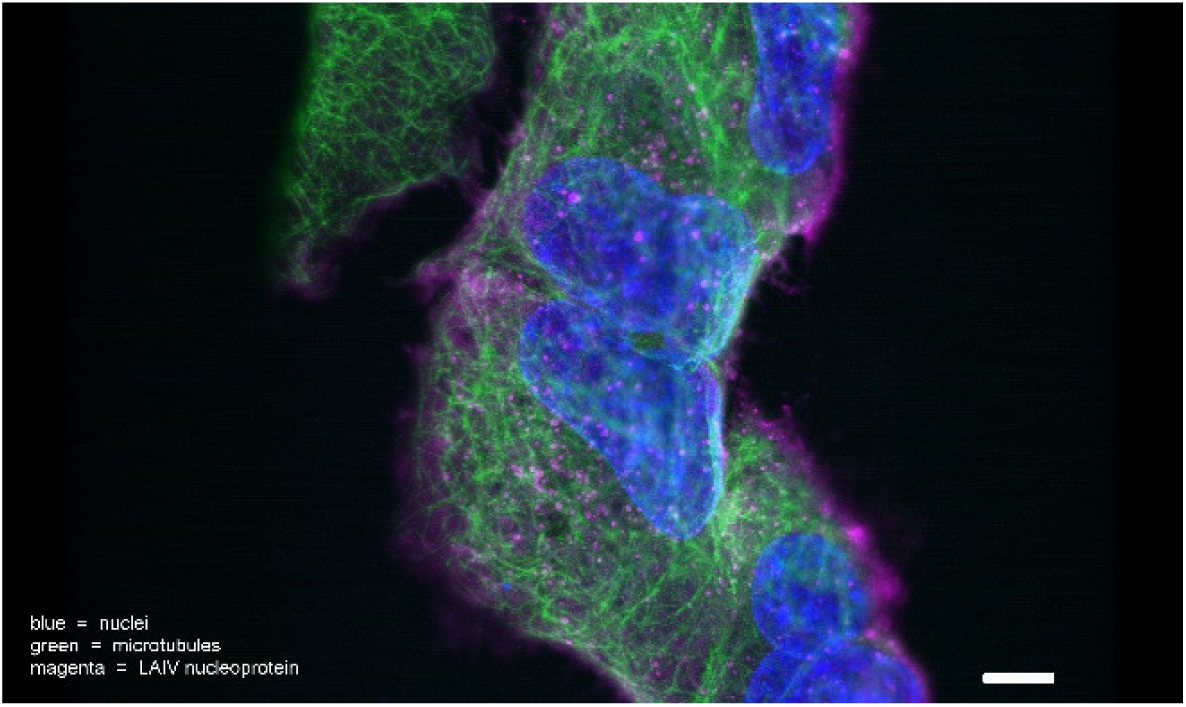
3D rotation of an expanded sample (A549 cells infected with live attenuated influenza vaccine) imaged on a light sheet microscope. Blue shows DAPI staining of the cell nucleus, green the cell microtubules and magenta the LAIV nucleoprotein (NP). Scale bar 20 µm.

